# Joint likelihood-free inference of the number of selected single nucleotide polymorphisms and the selection coefficient in an evolving population

**DOI:** 10.1101/2022.09.20.508756

**Authors:** Yuehao Xu, Andreas Futschik, Ritabrata Dutta

## Abstract

Given the often intractable exact likelihood, likelihood-free inference plays an important role in population genetics. Indeed, several methodological developments in approximate Bayesian Computation (ABC) were inspired by applications in population genetics. Here, we explore a novel combination of recently proposed ABC tools that can handle high-dimensional summary statistics and apply them to infer selection strength and the number of selected loci from experimental evolution data. While several methods infer selection strength at the single-nucleotide polymorphism (SNP) level, our approach provides additional information about the selective architecture, including the number of selected positions in a candidate window of interest. This is nontrivial, since the spatial correlation induced by genomic linkage can produce selection signals at neighboring SNPs. A further advantage of our approach is that we can readily quantify uncertainty using the ABC posterior. On both simulated and real data, we demonstrate promising performance. This suggests that our ABC variant could also be interesting in other applications.

## 1. Introduction

Bayesian methods have gained wide use in recent decades, supported by the growth of computational resources and algorithmic tools. Markov Chain Monte Carlo (MCMC) methods, in particular, allow sampling from posterior distributions even when analytic integration is intractable. Yet these methods still rely on evaluating the likelihood function. In many scientific applications, however, the likelihood itself is unavailable or prohibitively expensive to compute. This is especially true in population genetics, where dependencies across genomic loci arise from an unknown genealogical history. The large number of possible histories makes exact likelihood evaluations typically infeasible (Beaumont et al., 2002). Thus, likelihood-free methods are popular in this field, as they only require simulating the underlying model (Yuan et al., 2012). In recent years, considerable effort has been put into developing methodologies for likelihood-free inference (LFI) (Lintusaari et al., 2017). Approximate Bayesian Computation (ABC) (Beaumont et al., 2002; Pritchard et al., 1999) and synthetic likelihood (Wood, 2010) are among the most popular approaches, and several variants have been proposed that differ in terms of simulation strategies, as well as the choice of distance measures and summary statistics. In any case, proper tuning is necessary to optimize the performance in a given application. For a more complete review of LFI see Beaumont (2010a), Hartig et al. (2011), and Sisson et al. (2018).

One important application area of likelihood-free methods is the analysis of genomic time series data to detect signals of selection. In recent years, different tools have been proposed to search for such signals of selection. A composite-likelihood approach for evolve and resequence experiments (CLEAR) has been proposed by Iranmehr et al. (2017) to detect selected loci and estimate their selection coefficients. A Bayesian approach called WFABC (Foll et al., 2015) provides estimates of the effective population size (*N*_*e*_), selection and dominance coefficients under the Wright-Fisher model. A recent comparison of methods for detecting and quantifying selection based on data from evolve and resequence experiments, can be found in Vlachos et al. (2019).

Several statistical methods have also been proposed for the related but different challenge to detect the presence of selection in natural populations (e.g., (Tajima, 1983; Fay and Wu, 2000; Kim and Stephan, 2002; Kim and Nielsen, 2004; Nielsen et al., 2005; Foll and Gaggiotti, 2008; Pavlidis et al., 2013; Ferrer-Admetlla et al., 2014; DeGiorgio et al., 2016)). These methods rely on present-day data. Alternatively, historical samples are sometimes used. Past genomic information may include ancient DNA or derive from older samples preserved in Biobanks. Historic samples also contribute data from additional time points, but the underlying data-generating process is not controlled as in experimental evolution. In such a context, Malaspinas et al. (2012), He et al. (2020), and Schraiber et al. (2016) provide methods to infer selection along with allele ages from ancient genomic data based on Hidden Markov Models (HMM) using a diffusion approximation of the Wright-Fisher model. Further approaches in this direction are proposed by Lacerda and Seoighe (2014), Feder et al. (2014), Terhorst et al. (2015), and Paris et al. (2019). For natural populations, some authors have also proposed methods that jointly infer demography and selection. The methods by Sackman et al. (2019), Johri et al. (2020), and Fraïsse et al. (2021) utilize ABC for this purpose.

These traditional approaches to detecting selection often operate at the level of individual SNPs (single-nucleotide polymorphisms), typically assuming that only one site within a genomic region is under selection. While this assumption reduces computational complexity, it yields an overly sparse causal model that may not reflect biological reality, in which adaptation can result from the combined action of multiple linked loci. Ignoring this possibility may lead to misleading conclusions, such as overestimating the effect size of a single variant or overlooking more complex selective signals. To address this, we consider the number of selected loci *n*_*sel*_ and the corresponding selection coefficients 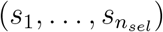 as jointly estimable parameters. The former quantifies the strength of selection, while the latter characterizes how that selection is distributed—whether concentrated at a single site or dispersed across multiple positions within the region. Estimating both parameters allows us to contextualize the intensity of selection within the local genetic architecture and distinguish different modes of adaptation. This formulation is related to problems in model selection, where one seeks not only to detect the presence of a signal but also to recover its underlying structure. By explicitly parameterizing both selection magnitude and locus-level configuration, we provide a model that goes beyond single locus inference to identify the number of selected positions within a window of interest. Notice that our model does permit the inference of locus-specific selection coefficients.

Our focus is on experimental evolution, where populations are exposed to a defined stressor under controlled conditions. Over several generations, allele frequencies change in response to selection. Whole-genome sequencing at multiple time points provides time-series data to monitor these changes. Replication is achieved by starting the experiment multiple times with genetically identical populations. Here, we propose an ABC framework that simultaneously estimates both the selection coefficient and the number of selected loci within genomic windows of interest. Such windows might be chosen based on functional insights or pre-screening for regions affected by selection. Our approach targets adaptation signals involving multiple linked sites rather than a single SNP. Our *main methodological novelty* lies in the choice of summary statistics and distance functions for ABC, specifically suitable for this task. For each SNP and time interval, we compute an approximate selection coefficient using the logit transformation described in Taus et al. (2017). These estimates, obtained independently across replicates that preserve both temporal and genomic structure, are used as summary statistics. Multiple samples of this summary statistics induce a distribution on function space, motivating us to use the expected energy score (Gneiting and Raftery, 2007) as a distance function, which is a metric on probability distributions on function space.

In section 2, we describe the discrete-time Wright-Fisher model used as the simulation model to generate synthetic data. The proposed inferential method based on ABC is described in Section 3, including details on the sampling scheme, summary statistics, and distances used in ABC. We study the performance of our proposed methodology on simulated datasets under different scenarios in Section 4. In Section 5, we apply them to an experimental Yeast dataset (Burke et al., 2014). We chose yeast as the organism in our real data example, since all founder haplotypes were available for this experiment. A discussion of our results and future directions in Section 6 concludes the manuscript.

## 2. Temporal Population Genetic Model

Since our proposed method is fundamentally simulation-based, we will use a simulator that generates data mimicking real-world observations, enabling comparison between the simulated and observed datasets. In Figure 1, we illustrate a typical evolve-and-resequence experiment (E&R) that is used to generate real-life population genetic data. To mimic a dataset like this, we consider a Wright-Fisher model with selection (Wright, 1931; Fisher, 1930) here.

**Figure 1:**
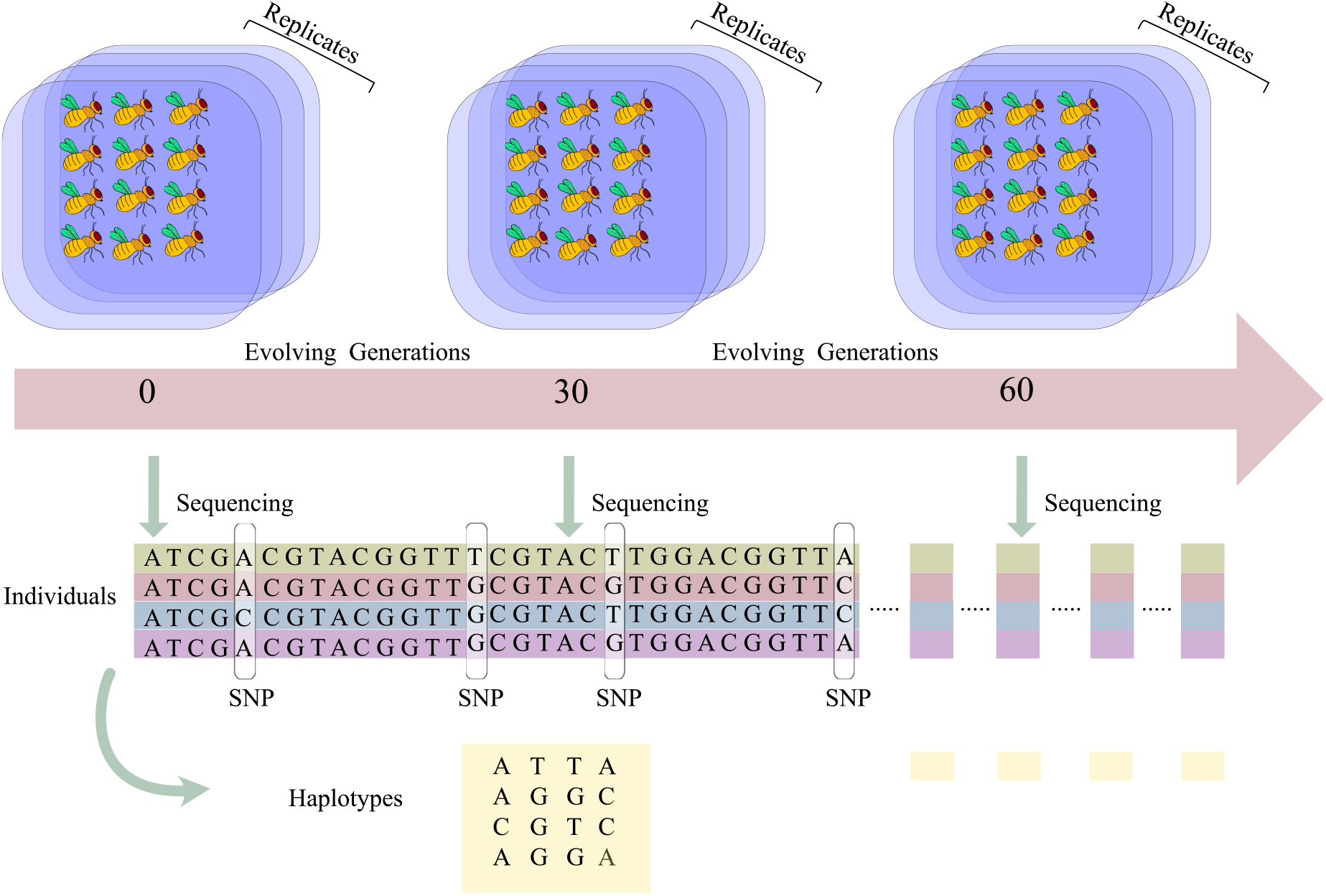
Design and data structure of a typical evolve-and-resequence experiment. Multiple replicate populations (blue) with controlled effective population size *N*_*e*_ evolve in the laboratory for 60 generations (pink). At the specified time points, a DNA pool is extracted from each replicate population and sequenced. This leads to estimated allele frequencies at each SNP. Time is measured in terms of generation numbers.

The simulations for these models could be carried out either in an exact discrete-time setup or in continuous time. Given our focus on precision, computational efficiency, and minimal bias, we have chosen the discrete-time Wright-Fisher model for our investigations. Existing software for forward-time simulation of such a model includes simuPOP (Peng and Kimmel, 2005), fwdpp (Thornton, 2014), MimiCREE2 (Vlachos and Kofler, 2018), and SLiM3 (Haller and Messer, 2019). In this work, we use the MimiCREE2 (Vlachos and Kofler, 2018) software to simulate from the discrete-time Wright-Fisher model and consider both haploid and diploid individuals.

### 2.1. Discrete time Wright-Fisher Model

Sewall Wright (Wright, 1931) and R.A. Fisher (Fisher, 1930) proposed the so-called Wright-Fisher model, which is used extensively as a standard population genetic model. It is characterized by discrete, non-overlapping generations and a constant population size, with the new generation drawn by random sampling with replacement from the gene pool of the previous generation. In practice, deviations from these assumptions of the Wright-Fisher model are often encountered. Therefore, the Wright-Fisher model is usually not applied with the census population size *N* but with the smaller effective population size *N*_*e*_ that produces the same amount of genetic drift as observed in the investigated population.

Under neutrality, the sampling probabilities under a Wright-Fisher model are given by the haplotype frequencies in the parental generation. Thus, expected frequencies stay constant across generations (Ewens, 2004). More generally, the haplotypes inherit the combined fitness of the SNPs they carry. For haploid individuals under selection, the probability of drawing a haplotype depends both on its previous frequency and its overall fitness. Assume that there are *m* SNPs with selection coefficients *s*_*i*_ (1 ≤ *i* ≤ *m*). Given *k* haplo-types (*h*_1_, *h*_2_, …, *h*_*k*_), the fitness of haplotype *h*_*j*_ will be *w*_*j*_ = 1 + Σ_*i*_ *s*_*i*_(*h*_*j*_). If haplotype *h*_*j*_ carries a neutral allele at position *i*, we have that *s*_*i*_(*h*_*j*_) = 0, and for beneficial and deleterious alleles we have *s*_*i*_(*h*_*j*_) *>* 0 and *s*_*i*_(*h*_*j*_) *<* 0 respectively. Assuming *N*_*Hap*_ haplotypes and a haploid organism, the haplo-type frequencies **f**_**t**+**1**_ in generation *t* + 1 are obtained via multinomial sampling using the frequencies **f**_**t**_ of the parental generation and the fitness vector **w** = (*w*_1_, …, *w*_*k*_). More specifically,

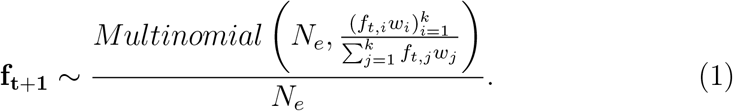

Neutrality corresponds to the special case where all *w*_*i*_ are equal.

In a diploid population, each individual carries two haplotypes. Under the additive model, the contribution of SNP *i* with two types of alleles *A* and *a* to the fitness of individual *J* is the sum of the contributions over both alleles carried by the individual: 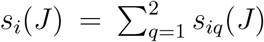. By specifying this additive model, we assume no dominance. A model that includes dominance would involve a more complex function than the sum, possibly depending on an unknown parameter, called the dominance coefficient. We do not consider dominance here, but note that its inclusion would complicate the inference problem. The overall contribution of a genomic region to the fitness of an individual is then obtained by summing over all SNPs within this region. In particular, for individual *J* we obtain the fitness *w*(*J*) = 1 + Σ_*i*_ *s*_*i*_(*J*). This assumption excludes epistasis, i.e. interaction effects between loci. To form the next generation, *N*_*e*_ individuals are randomly drawn with replacement and drawing probabilities proportional to their fitness. The chosen individuals contribute one of their segments, each with probability 1*/*2. In our simulation studies, we consider both the haploid and the diploid versions of the Wright-Fisher model.

The output of our simulator will be a matrix **x** = {*a*_*t,i*_ : *t* = *t*_0_, …, *T, i* = 1, …, *l*} where columns represent the SNPs, and rows represent the time points at which data are available. The entries are allele frequencies of the considered selected loci, resulting from the underlying haplotype frequencies {*h*_*t,i*_ : *t* = *t*_0_, …, *T, i* = 1, …, 2^*ℓ*^}. They are provided from the start time *t*_0_ to *T* = *t*_0_ +*K* × Δ*t*, at intervals of length Δ*t*. We thus assume that haplotype frequencies are observed at *K* + 1 equidistant time points. This assumption is not necessary for our method to work, provided that the simulated data are available at the same times as the observed data.

If we have *n* replicates in our observed data, we denote our observed data as 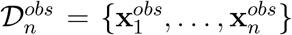. Similarly a simulated dataset with *m* replicates simulated using parameter *θ* would be denoted as 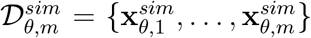. Here, we aim to jointly infer the number of selected SNPs (*n*_*sel*_) and the corresponding selection coefficients. We use independent Uniform(0, 0.2) distributions as priors for each selection coefficient, reflecting the absence of strong prior knowledge about their magnitudes in typical experimental evolution settings. This range covers weak to strong selection. A discrete uniform distribution with support 0, 1, 2 is considered as the prior for *n*_*sel*_. Given the maximum value of *n*_*sel*_ is two, we denote the selection coefficients at the first and second selected SNPs to be *s*_1_ and *s*_2_, and for *n*_*sel*_ = 1, we restrict *s*_2_ = 0 and *s*_1_ *>* 0. Hence, the considered parameters for our simulator model become *θ* = (*n*_*sel*_, *s*_1_, *s*_2_).

## 3. Likelihood-free inference of number of selected SNPs and selection coefficients

As in many population genetic models, the likelihood function is analytically intractable for the multivariate temporal population genetic model considered above. Nevertheless, the considered model allows direct sampling, a distinctive feature we capitalize on. This capability to directly sample from the model suggests using a likelihood-free inference framework. This approach entails generating data by running simulations based on the model parameters. The samples we generate from the simulator contain a spectrum of potential outcomes under the model, given the population parameters. By using methods available under the broad umbrella of likelihood-free inference (LFI), these samples let us make inferences without having to confront the mathematical intractability of the likelihood function (Beaumont et al., 2002; Foll et al., 2015; Collin et al., 2021). Here we use one of the most popular classes of LFI methods, approximate Bayesian computation (ABC) (Lintusaari et al., 2017), which approximates the intractable likelihood function implicitly using these samples. The asymptotic contraction of these approximate posteriors towards the true parameter value depends upon the choice of the summary statistics and some conditions being satisfied by the chosen summary statistics (Frazier et al., 2018; Li and Fearnhead, 2018; Frazier et al., 2023).

### Approximate Bayesian computation

The fundamental ABC rejection sampling scheme was first applied in the context of population genetics by Pritchard et al. (1999). Given observed dataset 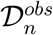, this iterates the following steps:

- Draw *θ* from the prior *π*(*θ*).
- Simulate a synthetic dataset 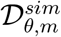 from the simulator-based model.
- Accept the parameter value *θ* if 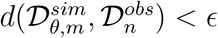. Otherwise, reject *θ*.

Here, the metric on the dataspace 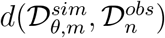 measures the closeness between 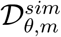 and 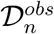. The accepted 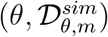 pairs are thus jointly sampled from a distribution proportional to 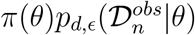, where 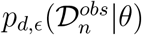 is an approximation to the likelihood function 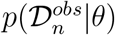:

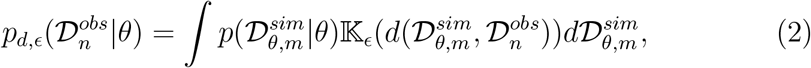

where 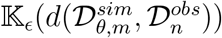 is in this case a probability density function proportional to 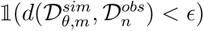^1^. Besides this choice for 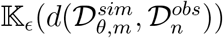, that has been exploited in several ABC algorithms (for instance Beaumont (2010b); Drovandi and Pettitt (2011); Del Moral et al. (2012); Lenormand et al. (2013)), ABC algorithms relying on different choices exist, for instance being proportional to 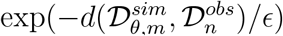 in simulated-annealing ABC (SABC) (Albert et al., 2015). In general, 𝕂_*ϵ*_(·) needs to be a probability density function with a large concentration of mass near 0, in which the parameter *ϵ* denotes the amount of concentration (the smaller *ϵ*, the more concentrated the density is). This guarantees that, in principle, the above approximate likelihood converges to the true one when *ϵ* → 0. Of course, lowering the threshold increases the computational cost, since fewer simulations are accepted.

More advanced algorithms than the simple rejection scheme detailed above are possible, for instance ones based on population or sequential Monte Carlo (Beaumont, 2010b; Del Moral et al., 2012; Lenormand et al., 2013), in which various parameter-data pairs are considered at a time and are evolved over several generations, while *ϵ* is decreased towards 0 at each generation to improve the approximation of the likelihood function, so that you are able to approximately sample from the true posterior distribution.

For the inference of parameters of our temporal population genetics model, we use the population Monte Carlo approximate Bayesian computation (PM-CABC) algorithm, proposed in Beaumont (2010b), for its theoretical rigour, few tuning parameters, and suitability for high-performance computing systems (Dutta et al., 2021). At the first step of this algorithm, *N*_sample_-many parameter values are randomly drawn from the prior distribution, and the value of *ϵ* is decreased by choosing the 50-th quantile of distances of the pseudo data simulated from the model using those randomly sampled parameter values in the previous step. In the next step, we produce *N*_sample_-many parameter values approximately distributed from the distribution *p*_*d,ϵ*_(*θ*|**x**^*obs*^), for the adapted *ϵ* value from the last step and again decrease the *ϵ* depending on the new samples. This procedure is repeated *N*_step_ times or until a stopping criterion is met. We note that the adapted *ϵ* values at each step are strictly decreasing and converge to zero, therefore improving the approximation to the posterior distribution. We use the algorithm using parallelized implementation available in *abcpy* Python package (Dutta et al., 2021).

### Choice of summary statistics

A common practice in ABC literature is to define *d* as the Euclidean distance between lower-dimensional summary statistics *S* : **x ↦** *S*(**x**), which, if sufficient, provide us with a consistent posterior approximation (Frazier et al., 2018; Li and Fearnhead, 2018). As sufficient summary statistics are not known for most of the complex models, the choice of summary statistics remains a problem (Csilléry et al., 2010), and they have been previously chosen in a problem-specific manner (Blum et al., 2013; Fearnhead and Prangle, 2012; Gutmann et al., 2018). Furthermore, they reduce the dimensionality of the original data while preserving as much information as possible.

Previously proposed estimates and test statistics for the selection coefficient (*s*) include those by Tajima (1983), Fay and Wu (2000), Kim and Stephan (2002), Kim and Nielsen (2004), Nielsen et al. (2005), Feder et al. (2014), Wiberg et al. (2017), Topa et al. (2015), Foll and Gaggiotti (2008), Pavlidis et al. (2013), Ferrer-Admetlla et al. (2014), and Kelly and Hughes (2019). Other quantities that are influenced by *s*, such as estimates of the effective population size (*N*_*e*_) have also been used as summary statistics before, see Foll et al. (2015). The estimation of *N*_*e*_ for both natural and experimental populations has been investigated by several authors (Waples (1989), Jorde and Ryman (2007), Jónás et al. (2016)). Here, we examined several potential summary statistics and found that estimates of *s* based on logit-transformed allele frequencies were most suitable. These summary statistics and the intuitions underlying them are explained below.

When *p*_*t*_ is the allele frequency at generation *t* ≥ 0, the allele frequency of a selected SNP in a haploid population can be modeled (Taus et al., 2017)

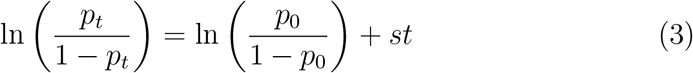

in the absence of drift and sampling noise. The analogous equation for the diploid case is

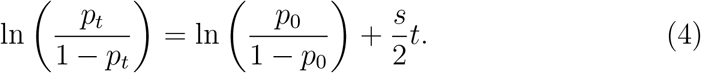

In the presence of drift, this formula remains a good approximation for inter-mediate allele frequencies when population and sample sizes are sufficiently large. (Therefore, we filter out SNPs with low and high starting frequencies.) Estimates of *s* are then computed by solving either equation (3) or (4) for all SNPs, and we sort them in ascending order by their values. An advantage of these estimates is that they are computationally efficient, thereby accelerating ABC computations.

Figures 2 and 3 provide examples that are based on one simulated data set for each considered scenario. They illustrate that our chosen summary statistics depend on both the selection strength and the number of loci (SNPs) under selection. Each data set has been generated using mimicrEE2 (Vlachos and Kofler, 2018). It involves ten replicate diploid populations with *N*_*e*_ = 1000, no recombination, and 60 generations. For 2 selected targets, the same selection strength (e.g., 0.02 or 0.07) is assumed for both. The heat maps in Figure 3 show that the estimates are more similar and lie around 0 when there is no selection (*n*_*sel*_ = 0). Under selection, larger estimates are present. These effects are more pronounced when *s* is large. The same patterns can also be seen in Figure 2, where the lines become steeper when the number of selected targets increases. The datasets used in the figures represent a single simulation run and are intended to illustrate typical patterns. Notice, however, especially for small values of *s*, there will be considerable random fluctuations across runs and the resulting patterns in summary statistics.

**Figure 2:**
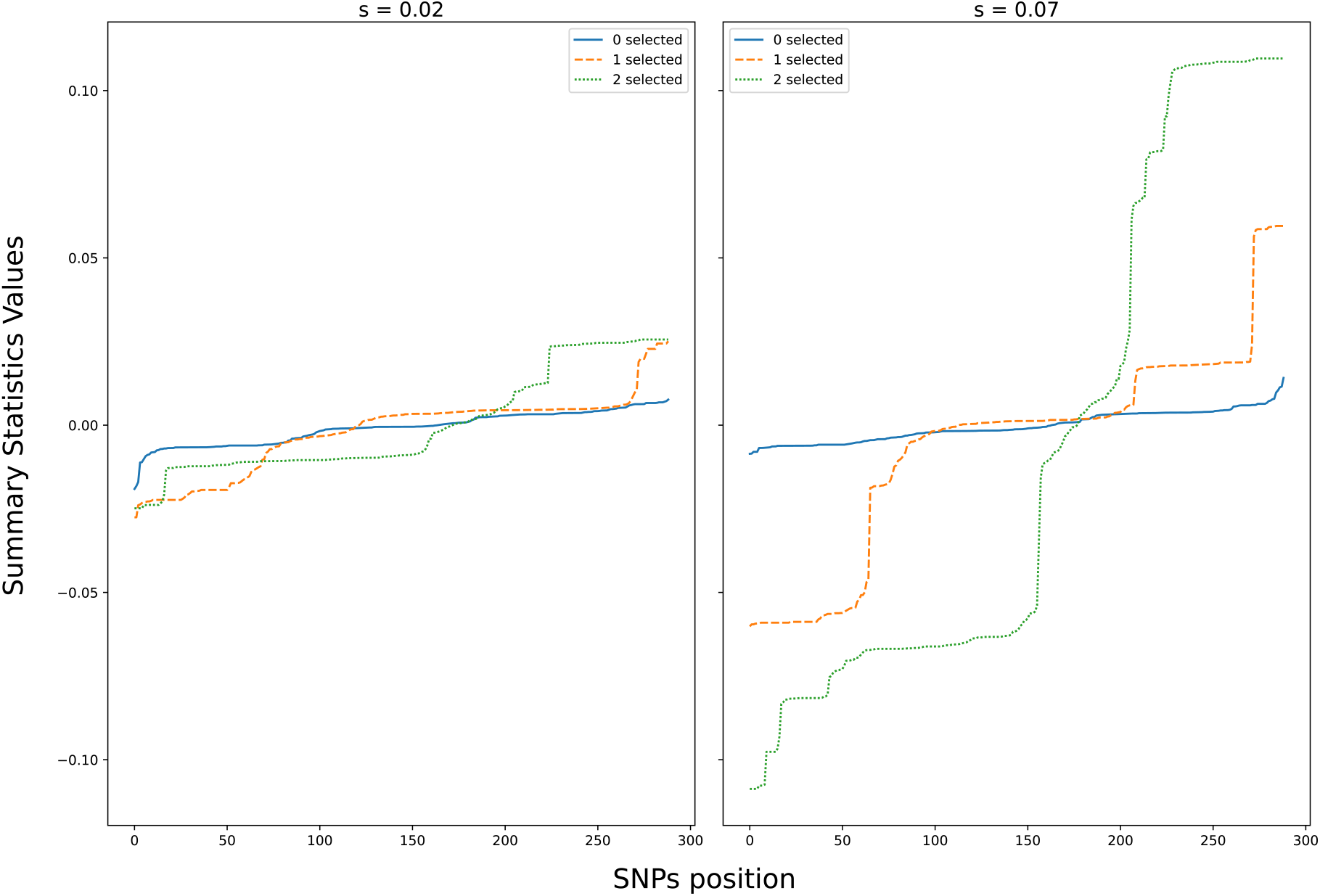
Average of the summary statistics over ten replicates for (*s*_1_, *s*_2_) = (0.02, 0.02) and (*s*_1_, *s*_2_) = (0.07, 0.07) from one simulation run. The Y-axis codes the values of summary statistics, while the X-axis represents the ordered SNP positions. Different colors represent different numbers of selected loci. The dataset consists of *N*_*e*_ = 1000 diploid individuals that evolve for 60 generations. For all SNPs, the corresponding estimates of *s* are computed from the allele frequencies at generations 0 and 60. For the scenario with a single selected SNP, the initial allele frequency is 0.191. For 2 selected targets, the starting allele frequencies are 0.183 and 0.187 respectively.

**Figure 3:**
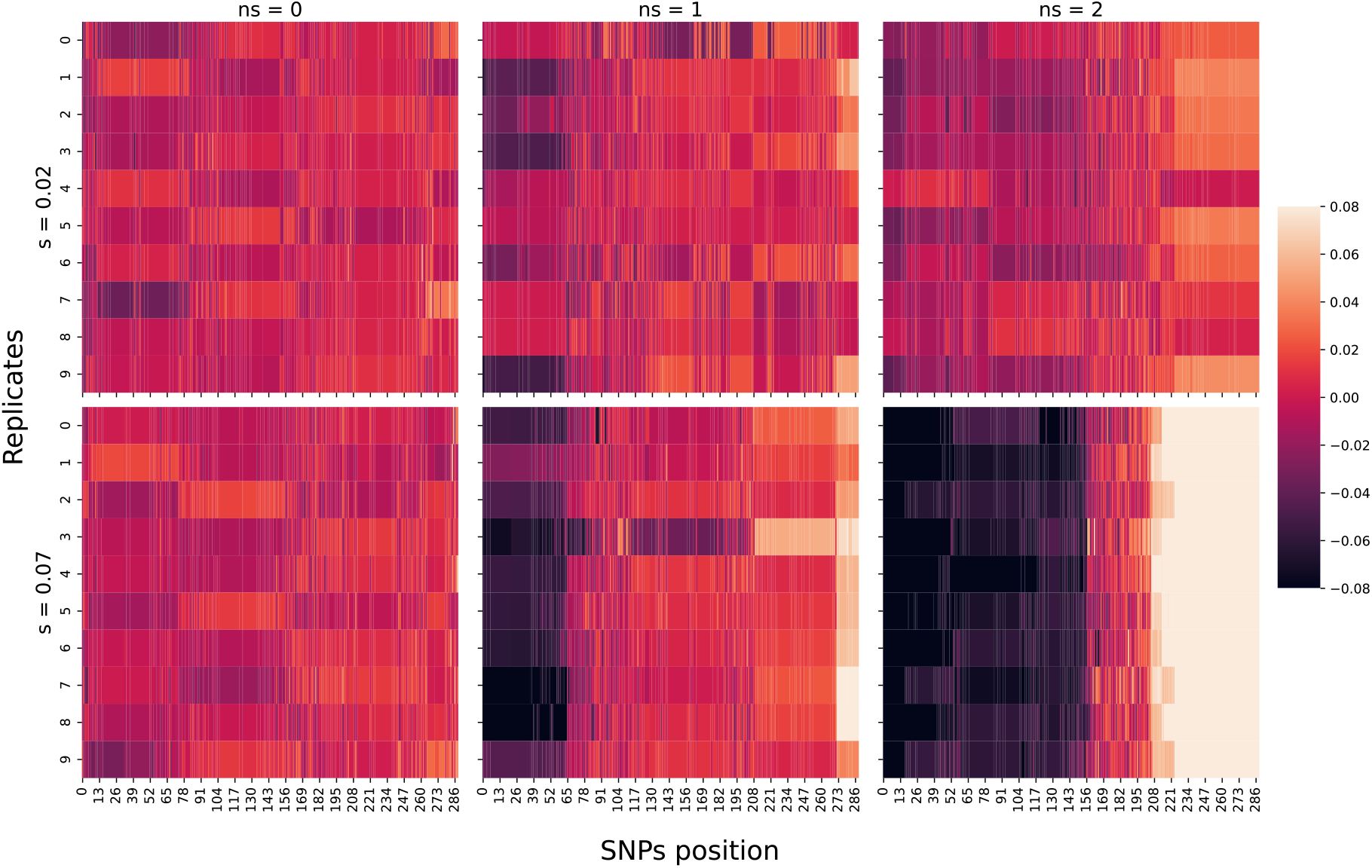
Heatmap of summary statistic values for different numbers of selected targets. The x-axis represents the position of the SNPs, and the y-axis represents replicates. The top maps are for *s* = 0.02, while the bottom maps are for *s* = 0.07. There are heatmaps for different numbers of selected targets (ns = 0 (left), 1 (center), 2(right column)).

### Expected energy score as distance

The extracted summary statistics *S*(**x**), a function of SNPs, when computed for replicates of observed dataset 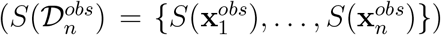 or simulated dataset 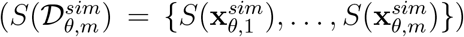, they provide us with samples from a distribution on function space. Hence, to compute 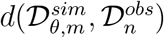, we consider the expected energy score (EES) (Gneiting and Raftery, 2007; Rizzo and Székely, 2016) as a distance between probability distributions on function space, which is a metric on probability distributions. Distances between probability measures have been used in ABC, when **x** is a set of independent and identically distributed (i.i.d.) draws (e.g., maximum mean discrepancy (Park et al., 2016), Energy score (Nguyen et al., 2020), Wasserstein distance (Bernton et al., 2019), Kullback-Leibler divergence (Jiang, 2018), classification accuracy estimating Jensen-Shannon Divergence (Gutmann et al., 2018), gamma-divergence (Fujisawa et al., 2021), and squared Hellinger distance (Frazier, 2020). The EES between two distributions *P* and *Q*, is defined as

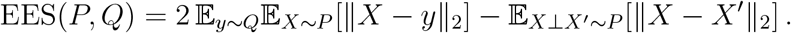

It is a strictly proper scoring rule for distributions satisfying 𝔼_*X*~*P*_ ∥*X*∥ *<* ∞ and induces a statistical divergence (Dawid, 2007; Pacchiardi et al., 2024). Given correspondingly *m* and *n* independent and identically distributed samples from *P* and *Q*, the expected energy score can be unbiasedly estimated as,

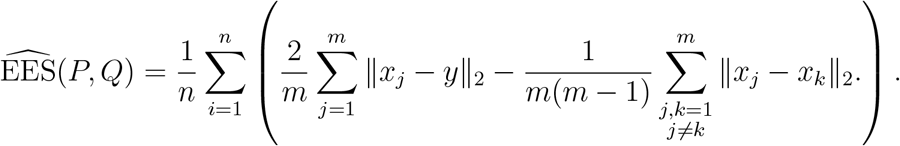

We further note that, for smaller values of *n* (the number of replicates present in observed experimental datasets in temporal population genetics could very well be fewer than 5), the above expression unbiasedly estimates the energy scoring (*ES*) rule,

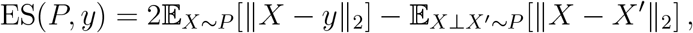

which provides a penalty when stating a distribution *P* for observation *y*. This has recently been used as a replacement for the log-likelihood in the framework of generalized Bayesian inference Pacchiardi et al. (2024). Using simulated and observed datasets, our proposed distance becomes:

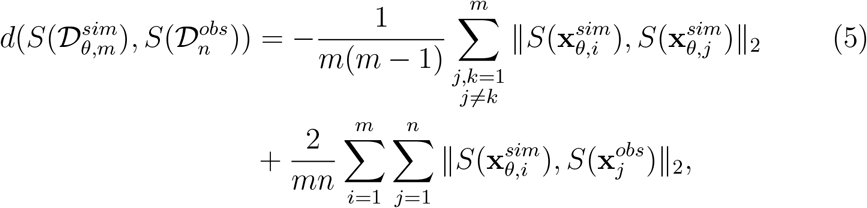

which can be interpreted as an unbiased estimate of the expected energy score, or the energy score itself, for large or small values of *n*.

### Estimation of parameters

The sampling scheme proposed here, using the summary and distance described before, provides us *N*_sample_-many 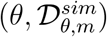 pairs from a distribution proportional to 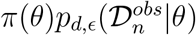. Marginalizing 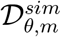, we get samples of *θ* = (*n*_*sel*_, *s*_1_, *s*_2_) from an approximate posterior distribution of (*n*_*sel*_, *s*_1_, *s*_2_) given observed data 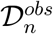,

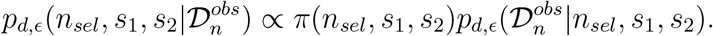

Our estimate of number of selected locations, 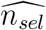 is the mode of the marginal approximate posterior distribution 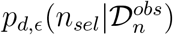. We estimate the selection coefficients as the mode of the conditional marginal distribution of (*s*_1_, *s*_2_) given 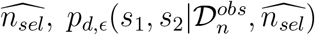. The mode is calculated via optimizing an estimated density of the conditional marginal distribution using a kernel density estimator.

## 4. Simulation Studies

We evaluate the performance of our approach by running inference over 20 simulated datasets, 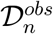, generated under varying biological scenarios. In particular, we consider different selection strengths (*s*_1_, *s*_2_), population sizes (*N*_*e*_), replicate numbers (*n* in observed dataset 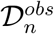). We also simulate both haploid and diploid organisms, diploids also with different recombination rates. The simulated data were generated under a discrete Wright-Fisher model using the MimiCREE2 (Vlachos and Kofler, 2018) simulator.

The software allows simulation with arbitrary founder sequences. Here, we use 189 founder haplotypes from a *Drosophila simulans* population sampled in Florida (Barghi et al., 2019). To make the simulations more comparable between the haploid and diploid scenarios, we used the same founder haplotypes in both simulations.

More specifically, our focus is on a window spanning positions 363-49337 on chromosome 2L. This window contains a total of 500 SNPs. In the preprocessing step, we filter out SNPs with starting allele frequencies below 0.1 or above 0.9, as their relative changes in allele frequency are more strongly affected by genetic drift and thus noisier. Furthermore, our chosen summary statistics require excluding SNPs that experience fixation or loss due to drift. Starting with very low or very high frequencies increases the likelihood of such events. The remaining 289 SNPs are used in the analysis.

They serve as the genetic input for both diploid and haploid scenarios. When simulating scenarios with positive selection, selected positions have been picked randomly among SNPs with starting allele frequencies between 0.1 and 0.5, as there is reasonable power to detect a signal from such SNPs. When an allele is already dominating in a population, there is not much room for a further allele frequency increase under positive selection. On the other hand, negative selection may be more easily detected and could indeed be identified as positive selection of the minor allele.

We simulate 60 generations, a timescale frequently adopted in laboratory selection experiments with higher organisms. It is long enough for selection signals to emerge and be detectable, yet short enough to retain polymorphism and avoid saturation.

Our inference methodology was run for each dataset, via PMCABC for 6 steps (= *N*_*step*_), generating 100 (= *N*_*sample*_) samples using 100 (= *m*) replicates for each simulated dataset 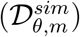. The mean runtime for each dataset was ~4 hours when all simulations were run on Dell PowerEdge C6420 compute nodes, each with 2 x Intel Xeon Platinum 8268 (Cascade Lake) 2.9 GHz 24-core processors, 48 cores per node, and 4 GB of RAM per core.

### Inference with haploid populations

In the haploid case, we first consider three scenarios with *n*_*sel*_ = 0, 1, 2, and, for each scenario, we vary the selection coefficients (*s*_1_, *s*_2_). In all of these cases, we consider 20 replicates per dataset and *N*_*e*_ = 1000. All the selection strengths considered, and the estimated 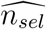 ans 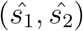 are reported correspondingly in Table 1 and Table 2. There, we also study a scenario with a smaller *N*_*e*_ = 500 under 2 selected locations. As shown in Table 1, selection is mostly detected when present. The stronger the selection, the higher the accuracy. This applies also to distinguishing between the cases of one and two selected loci. The table also provides an idea of how large both selection coefficients as well as the effective population size need to be for reliable identification of selection.

**Table 1:** Inference with haploid population. The estimated number of selected loci 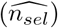 for 20 different datasets under different scenarios of varying the number of selected loci (*n*_*sel*_ = 0, 1, 2), selection strength, population size (*N*_*e*_ = 500, 1000), with 20 replicates in each observed dataset.

| #selected SNPs | $N_e$ | Selection strength | $\widehat{n_{sel}}$ over 20 experiments | | |
| --- | --- | --- | --- | --- | --- |
| | | | $\widehat{n_{sel}} = 0$ | $\widehat{n_{sel}} = 1$ | $\widehat{n_{sel}} = 2$ |
| $n_{sel} = 0$ | 1000 | $s : (0.0, 0.0)$ | 20 | 0 | 0 |
| $n_{sel} = 1$ | 1000 | $s : (0.02, 0.0)$ | 3 | 17 | 0 |
| | 1000 | $s : (0.05, 0.0)$ | 1 | 18 | 1 |
| | 1000 | $s : (0.07, 0.0)$ | 0 | 18 | 2 |
| $n_{sel} = 2$ | 1000 | $s : (0.05, 0.02)$ | 0 | 18 | 2 |
| | 1000 | $s : (0.05, 0.07)$ | 1 | 11 | 8 |
| | 1000 | $s : (0.07, 0.09)$ | 0 | 9 | 11 |
| $n_{sel} = 2$ | 500 | $s : (0.05, 0.02)$ | 0 | 13 | 7 |
| | 500 | $s : (0.05, 0.07)$ | 0 | 7 | 13 |
| | 500 | $s : (0.07, 0.09)$ | 0 | 6 | 14 |

**Table 2:** Inference with haploid population. The conditionally estimated selection coefficients 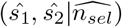 for 20 different datasets under different scenarios of varying number selected loci (*n*_*sel*_ = 0, 1, 2), selection strength, population size (*N*_*e*_ = 500, 1000), with 20 replicates in each observed dataset.

| #selected SNPs | $N_e$ | Selection strength | Mean / Variance of $\hat{s}$ over 20 experiments | |
| --- | --- | --- | --- | --- |
| | | | $\hat{s} \vert \widehat{n}_{sel} = 1$ | $\hat{s} \vert \widehat{n}_{sel} = 2$ |
| $n_{sel} = 0$ | 1000 | s : (0.0, 0.0) | - | - |
| $n_{sel} = 1$ | 1000 | s : (0.02, 0.0) | (0.018, 0) / (1.145e-5, 0) | - |
|  | 1000 | s : (0.05, 0.0) | (0.044, 0) / (0.00013, 0) | (0.041, 0.039) / (0, 0) |
|  | 1000 | s : (0.07, 0.0) | (0.062, 0.0) / (0.00019, 0.0) | (0.044, 0.07) / (0.00028, 5e-05) |
| $n_{sel} = 2$ | 1000 | s : (0.05, 0.02) | (0.041, 0.0) / (0.00036, 0.0) | (0.07, 0.041) / (0.00013, 0.00037) |
|  | 1000 | s : (0.05, 0.07) | (0.092, 0.0) / (0.00059, 0.0) | (0.072, 0.069) / (0.00144, 0.00034) |
|  | 1000 | s : (0.07, 0.09) | (0.078, 0.0) / (0.00062, 0.0) | (0.098, 0.069) / (0.00119, 0.00064) |
| $n_{sel} = 2$ | 500 | s : (0.05, 0.02) | (0.059, 0.0) / (0.00078, 0.0) | (0.054, 0.052) / (0.00089, 0.00049) |
|  | 500 | s : (0.05, 0.07) | (0.082, 0.0) / (0.00038, 0.0) | (0.072, 0.073) / (0.00095, 0.00152) |
|  | 500 | s : (0.07, 0.09) | (0.089, 0.0) / (0.00245, 0.0) | (0.079, 0.118) / (0.00077, 0.0009) |

According to Table 2, the estimates of the selection coefficients are reliable, given that the right model has been selected. The results suggest that even if there are two different loci, their selection coefficients can be reliably estimated given that the right model is chosen. If the number of selected loci is estimated incorrectly, the estimated selection coefficients typically do not reflect the true coefficients. It should be noted, however, that in view of Table 1, model misspecification is infrequent under sufficiently strong selection and occurs typically, if the observed data are atypical under the true model.

Due to cost constraints, actual evolutionary genetics experiments usually involve only 10-20 replicates or fewer. These replicate counts are commonly used in evolve-and-resequence studies, where constraints on cost and logistics limit feasibility. Increasing the number of replicates, for example, to 100, would enhance signal strength and facilitate inference, but this is often infeasible in real-world settings. To assess the effect of the replicate number, we assess their performance under two selected loci for 5, 10, and 20 replicates, and report the estimated 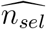 and 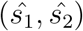 correspondingly in Table 3 and Table 4. It turns out that a reliable detection of *s*_*sel*_ using our method requires a sufficient number of selected loci. Clearly, five replicates are not enough to distinguish between one and two selected loci. Detecting selection altogether also seems to work with fewer replicates. Given that the correct model has been chosen, Table 4 suggests that the estimates of the selection coefficients are reliable for all chosen replicate numbers.

**Table 3:** Inference with haploid population - varying number of replicates. The estimated number of selected loci 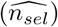 for 20 different datasets under different scenarios of varying selection strength and different replicates (5, 10, 20) in each observed dataset, under *n*_*sel*_ = 2 and fixed population size (*N*_*e*_ = 1000).

| #selected SNPs | replicates | Selection strength | Estimate of $n_{sel}$ over 20 experiments | | |
| --- | --- | --- | --- | --- | --- |
| | | | $\widehat{n}_{sel} = 0$ | $\widehat{n}_{sel} = 1$ | $\widehat{n}_{sel} = 2$ |
| $n_{sel} = 2$ | 5 | $s : (0.07, 0.09)$ | 0 | 14 | 6 |
| | 10 | $s : (0.07, 0.09)$ | 0 | 9 | 11 |
| | 20 | $s : (0.07, 0.09)$ | 0 | 9 | 11 |

**Table 4:**
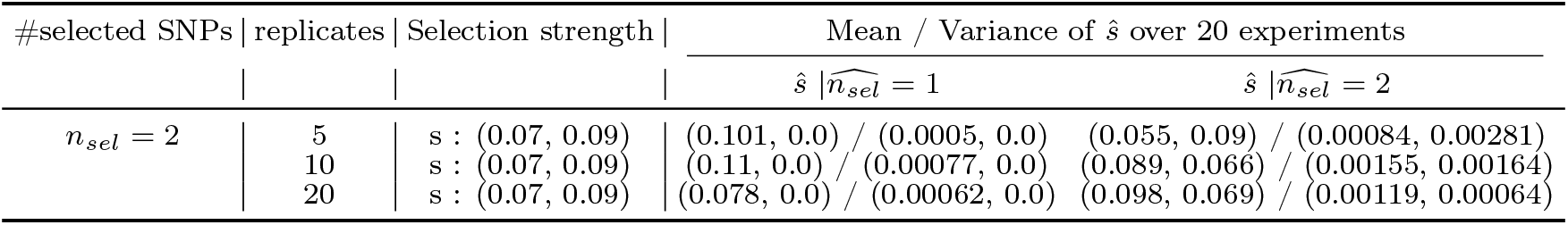
Inference with haploid population - varying number of replicates. The conditionally estimated selection coefficients 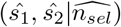 for 20 different datasets under different scenarios of varying selection strength and different replicates (5, 10, 20) in each observed dataset, under *n*_*sel*_ = 2 and fixed population size (*N*_*e*_ = 1000).

### Inference with diploid population

Next, in the diploid case, we consider three scenarios with *n*_*sel*_ = 0, 1, 2, and for each scenario, we vary the selection coefficients (*s*_1_, *s*_2_). In all of these cases, we consider 20 replicates per dataset and *N*_*e*_ = 1000. All the selection strengths considered, and the estimated 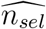 ans 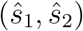 are reported correspondingly in Table 5 and Table 6. There, we also study a scenario with a smaller *N*_*e*_ = 500 under 2 selected loci. Because recombination is possible in a diploid population, we consider a recombination rate of 2.5 cM/Mb, a common value for *Drosophila simulans*. To better understand the effect of recombination, we also consider recombination rates 0, 2.5, 15, 20, and 49 cM/Mb in Tables 7 and 8.

**Table 5:** Inference with diploid population. The estimated number of selected loci 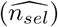 for 20 different datasets under different scenarios of varying number selected loci (*n*_*sel*_ = 0, 1, 2), selection strength, population size (*N*_*e*_ = 500, 1000) under fixed recombination rate of 2.5 cM/Mb and 20 replicates in each observed dataset.

| #selected SNPs | $N_e$ | Selection strength | Estimate of $n_{sel}$ over 20 experiments | | |
| --- | --- | --- | --- | --- | --- |
| | | | $\widehat{n}_{sel} = 0$ | $\widehat{n}_{sel} = 1$ | $\widehat{n}_{sel} = 2$ |
| $n_{sel} = 0$ (recomb. rate: 0) | 1000 | s : (0.0, 0.0) | 20 | 0 | 0 |
| $n_{sel} = 1$ (recomb. rate: 0) | 1000 | s : (0.07, 0.0) | 6 | 13 | 1 |
|  | 1000 | s : (0.1, 0.0) | 4 | 14 | 2 |
|  | 1000 | s : (0.14, 0.0) | 3 | 15 | 2 |
|  | 1000 | s : (0.18, 0.0) | 1 | 2 | 17 |
| $n_{sel} = 2$ (recomb. rate: 2.5) | 1000 | s : (0.05, 0.05) | 4 | 16 | 0 |
|  | 1000 | s : (0.07, 0.09) | 0 | 19 | 1 |
|  | 1000 | s : (0.09, 0.12) | 1 | 16 | 3 |
|  | 1000 | s : (0.14, 0.18) | 1 | 6 | 13 |
| $n_{sel} = 2$ (recomb. rate: 2.5) | 500 | s : (0.05, 0.05) | 2 | 1 | 17 |
|  | 500 | s : (0.07, 0.09) | 1 | 15 | 4 |
|  | 500 | s : (0.09, 0.12) | 1 | 10 | 9 |
|  | 500 | s : (0.14, 0.18) | 3 | 7 | 10 |

**Table 6:**
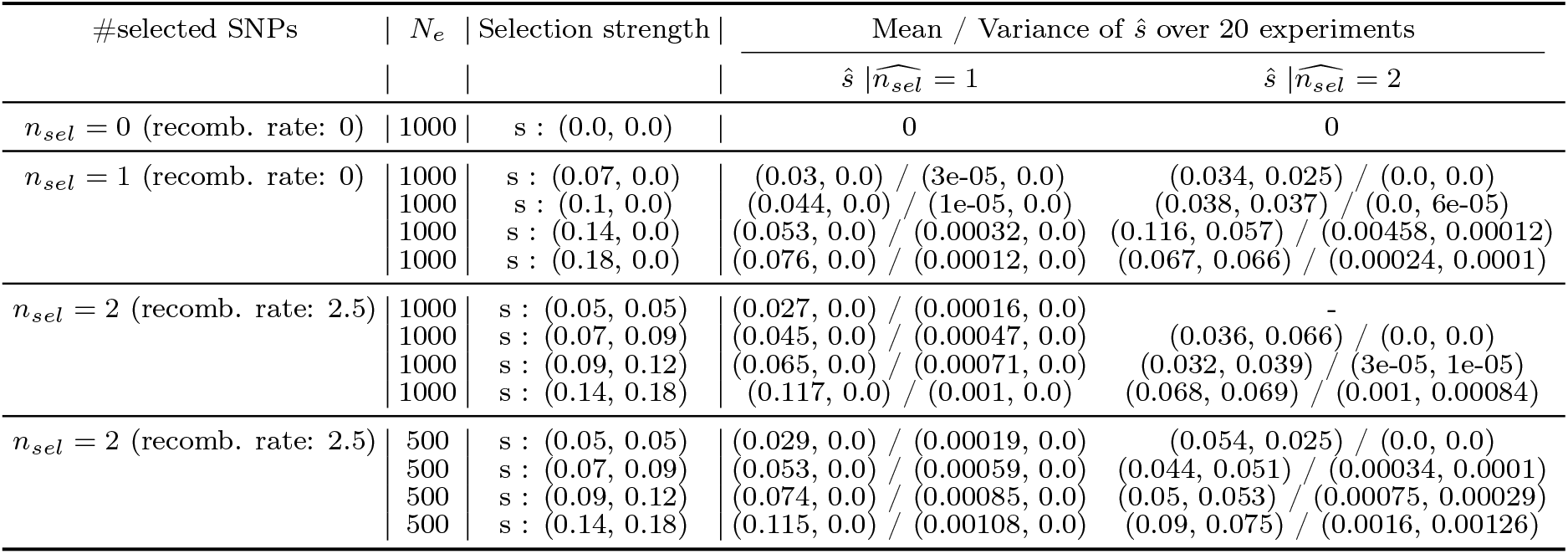
Inference with diploid population. The conditionally estimated selection coefficients 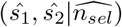 for 20 different datasets under different scenarios of varying number selected loci (*n*_*sel*_ = 0, 1, 2), selection strength, population size (*N*_*e*_ = 500, 1000) under fixed recombination rate of 2.5 cM/Mb and 20 replicates in each observed dataset.

**Table 7:** Inference with diploid population - varying recombination rates. The estimated number of selected loci 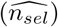 for 20 different datasets under different recombination rates (0, 2.5, 15, 30, 49 cM/Mb), under *n*_*sel*_ = 2 (*s*_1_, *s*_2_) = (0.14, 0.18) and fixed population size (*N*_*e*_ = 1000).

| #selected SNPs | recomb. rate | Selection strength | Estimate of $nns$ over 20 experiments | | |
| --- | --- | --- | --- | --- | --- |
| | | | $\widehat{n}_{sel} = 0$ | $\widehat{n}_{sel} = 1$ | $\widehat{n}_{sel} = 2$ |
| $n_{sel} = 2$ | 0 | $s : (0.14, 0.18)$ | 0 | 11 | 9 |
| | 2.5 | $s : (0.14, 0.18)$ | 3 | 7 | 10 |
| | 15 | $s : (0.14, 0.18)$ | 0 | 8 | 12 |
| | 30 | $s : (0.14, 0.18)$ | 1 | 9 | 10 |
| | 49 | $s : (0.14, 0.18)$ | 0 | 13 | 7 |

**Table 8:** Inference with diploid population - varying recombination rates. The conditionally estimated selection coefficients 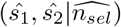 for 20 different datasets under different recombination rates (0, 2.5, 15, 30, 49 cM/Mb), under *n*_*sel*_ = 2, (*s*_1_, *s*_2_) = (0.14, 0.18) and fixed population size (*N*_*e*_ = 1000).

| #selected SNPs | recomb. rate | Selection strength | Mean / Variance of $\hat{s}$ over 20 experiments | |
| --- | --- | --- | --- | --- |
| | | | $\hat{s} \vert \widehat{n}_{sel} = 1$ | $\hat{s} \vert \widehat{n}_{sel} = 2$ |
| $n_{sel} = 2$ | 0 | $s : (0.14, 0.18)$ | (0.084, 0.0) / (0.00142, 0.0) | (0.054, 0.053) / (0.0004, 7e-05) |
| | 2.5 | $s : (0.14, 0.18)$ | (0.117, 0.0) / (0.001, 0.0) | (0.068, 0.069) / (0.001, 0.00084) |
| | 15 | $s : (0.14, 0.18)$ | (0.111, 0.0) / (0.00072, 0.0) | (0.074, 0.072) / (0.00132, 0.00091) |
| | 30 | $s : (0.14, 0.18)$ | (0.097, 0.0) / (0.00158, 0.0) | (0.072, 0.067) / (0.00129, 0.00054) |
| | 49 | $s : (0.14, 0.18)$ | (0.098, 0.0) / (0.0014, 0.0) | (0.065, 0.061) / (0.00092, 0.00025) |

For comparison with the haploid case, note that the selection coefficients are specified for homozygote individuals in mimiCREE2. This means that a selection coefficient of 2*s* is required to achieve the same strength of selection as in the haplod case. Thus, inference on the number of selected loci requires sufficiently strong selection in the diploid case. The estimated selection coefficients appear to be substantially underestimated in some scenarios.

## 5. Inference on Yeast dataset (Burke et al., 2014)

Here, we illustrate the application of our method to real data, specifically the outcrossing yeast data from Burke et al. (2014). The outcrossing yeast populations used in this study were initially derived from a cross of four founder strains: a wine strain (European), a West African strain, a North American strain, and a sake strain (Asian). Over 18 weeks, each population underwent forced outcrossing and periodic sampling, creating a unique sexual yeast model for studying adaptation through standing genetic variation. This structure enabled the investigation of evolutionary dynamics across multiple generations, making it particularly suitable for methods such as ours.

Previous yeast studies (Burke et al., 2014; Iranmehr et al., 2017; Taus et al., 2017) identified specific genomic regions, including chromosomes 9 and 11, where genome-wide scans detected significant signals of selection. These findings indicated that evolutionary changes were repeatable in response to experimental selection pressures across replicate populations. By focusing on these identified regions, we aim to validate the effectiveness of our method in detecting and characterizing selection effects within specific genomic intervals. Borrowing from those studies, we choose chromosome 11 and consider a window of 1000 SNPs (positions from 3049 to 168836). Chromosome 11 was selected for analysis because allele frequency shifts in this region were observed across multiple generations, suggesting adaptive selection. The data set contains allele frequencies at four time points (generations {0,180,360,540}) from 12 replicate populations.

Since the original study did not specify exact selection coefficients, the results from existing methods by Iranmehr et al. (2017) were used as reference points to validate findings. In Iranmehr et al. (2017), selection appears weak, showing the largest estimate of *s* was 0.009. Additionally, they also provide the estimated 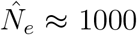 for the effective population size. In our simulator model, we use the estimated value of *N*_*e*_ and the starting haplotypes provided by Burke et al. (2014) to simulate allele frequencies. We note that simulating allele frequencies over 540 generations at 1000 SNPs is computationally expensive, making the direct application of our methodology infeasible across the entire region. Further, in the simulation studies, we investigated the behaviour of *n*_*sel*_ estimation for values in {0, 1, 2}. In the whole region, the true value of *n*_*sel*_ may exceed this range across the entire region. To make our inference computationally feasible and trustworthy, we split the region into 7 windows (the first 6 containing 140 SNPs and the last one containing 160 SNPs), run our inference scheme for each of these windows in parallel.

Assuming our yeast populations are haploid, we used the MimiCREE2 (Vlachos and Kofler, 2018) software as a simulator for inference. Our inference methodology was run for each window, where PMCABC was run for 5 steps, generating 50 (= *N*_*sample*_) samples using 50 (= *m*) replicates for each simulated dataset 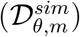. The mean runtime for each window was ~6 hours, and all runs were performed in parallel on Dell PowerEdge C6420 compute nodes, each with 2 x Intel Xeon Platinum 8268 (Cascade Lake) 2.9 GHz 24-core processors, 48 cores per node, and 4 GB of RAM per core. Given that high-performance computing facilities now have access to 1000s of similar nodes or better, this illustration also provides a straightforward pathway to scale this method to genome-wide studies.

Inference considering the 12 replicates concludes that there is no selection in any of the windows (Table 9), which is very similar to the results of Iranmehr et al. (2017), indicating selection strength of atmost 0.009. Further investigation of the summary statistics shows that 2 of the 12 replicates in the dataset exhibit strong selection effects (see Figure 4). In related work, genetic redundancy has often been mentioned as a possible reason for an inconsistency in the signal between replicates. This basically means that different replicates follow different genetic paths towards adaptation. Therefore, we rerun our inference using only the 2 informative replicates for the 7 windows, which concludes a significant number of selected SNPs in the first 4 windows, reported in Table 10 and Figure 5.

**Table 9:** Inference on Yeast data (using 12 replicates) is illustrated by reporting the estimated posterior probability of *nns*, inferred using all 12 replicates from the experimental data reported in Burke et al. (2014), assuming population size (*N*_*e*_ = 1000), for 7 windows on chromosome 11 each containing 140/160 SNPs.

| Windows | Positions (#SNPs) | $P(n_{sel} = 0\vert \mathcal{D}^{obs})$ | $P(n_{sel} = 1\vert \mathcal{D}^{obs})$ | $P(n_{sel} = 2\vert \mathcal{D}^{obs})$ |
| --- | --- | --- | --- | --- |
| 1 | 3049–37885 (140) | 0.84 | 0.16 | 0.0 |
| 2 | 37893–55213 (140) | 0.95 | 0.05 | 0.0 |
| 3 | 55421–76297 (140) | 0.98 | 0.12 | 0.0 |
| 4 | 76368–94611 (140) | 1.0 | 0.0 | 0.0 |
| 5 | 94682–115512 (140) | 1.0 | 0.0 | 0.0 |
| 6 | 115590–132426 (140) | 1.0 | 0.0 | 0.0 |
| 7 | 132468–168836 (160) | 1.0 | 0.0 | 0.0 |

**Figure 4:**
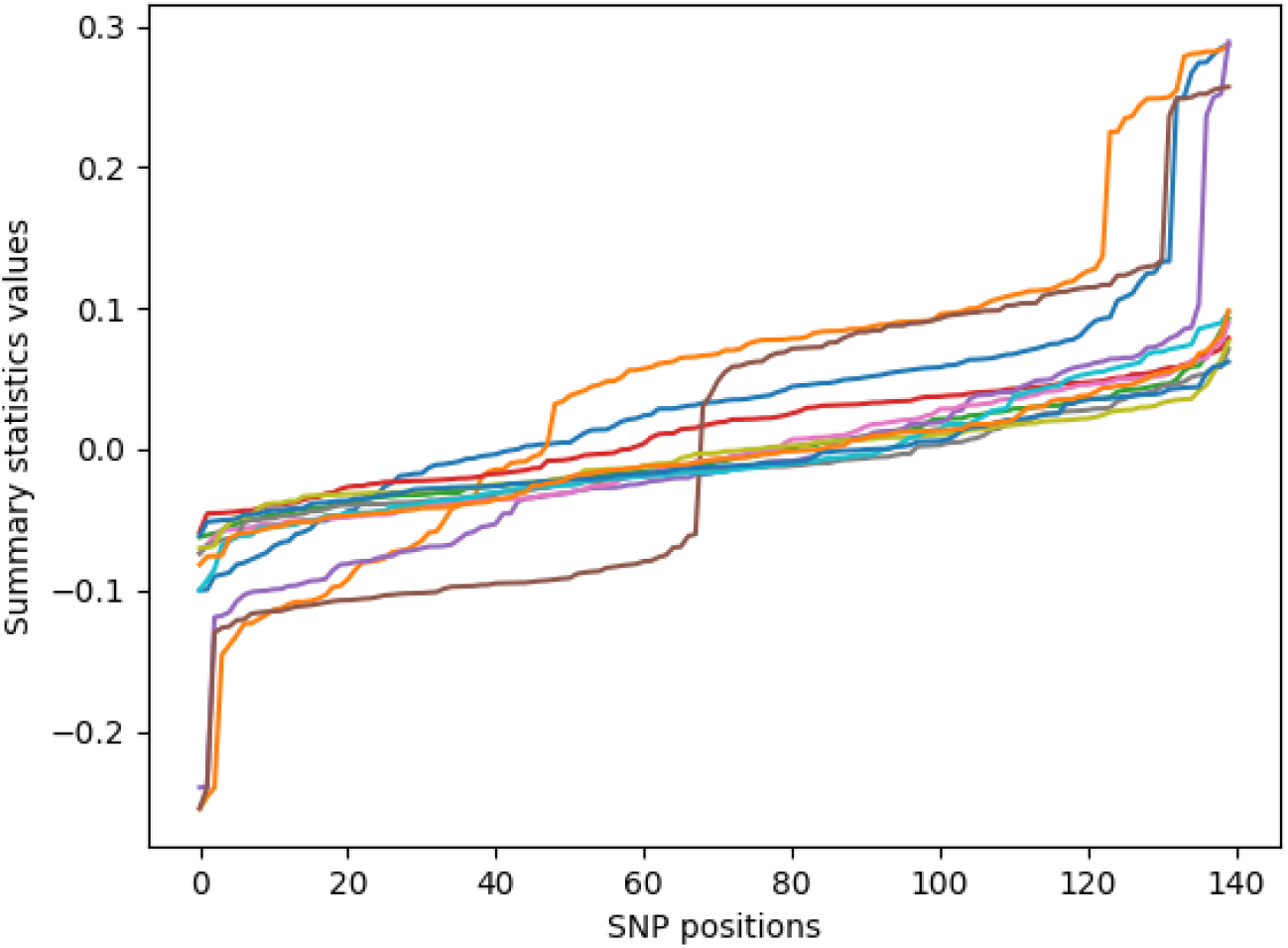
Summary statistics for Yeast data. We illustrate the calculated summary statistics for the 12 replicates (different colours stand for different replicates) of a Yeast dataset. We consider subwindow 1 from our considered region spanning positions 3049– 37885 on chromosome 11, which shows only two replicates with significant selection signals.

**Table 10:**
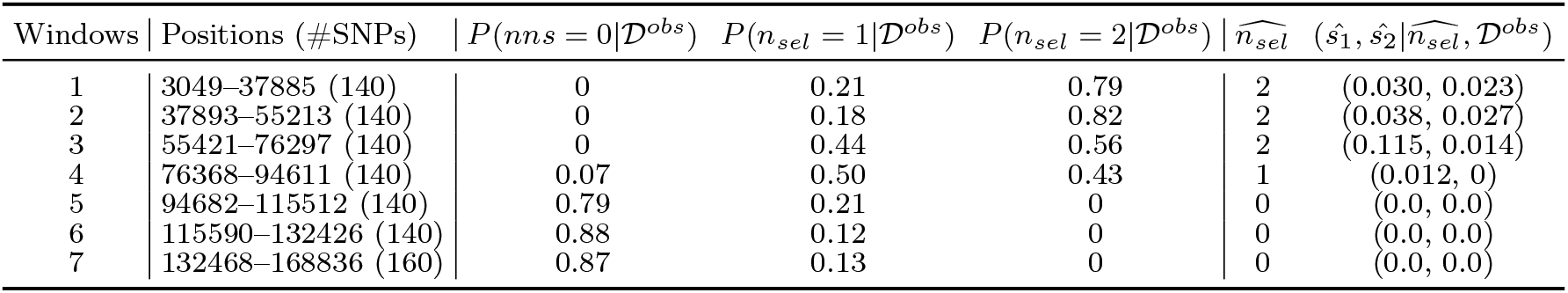
Inference on Yeast data (using 2 informative replicates) is illustrated by reporting the estimated posterior probability of *n*_*sel*_, estimated number of selected loci 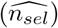 and the conditionally estimated selection coefficients 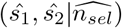 inferred using 2 replicates from the experimental data reported in Burke et al. (2014), assuming population size (*N*_*e*_ = 1000), for 7 windows on chromosome 11 each containing 140/160 SNPs.

| Windows | Positions (#SNPs) | $P(nns = 0\vert \mathcal{D}^{obs})$ | $P(n_{sel} = 1\vert \mathcal{D}^{obs})$ | $P(n_{sel} = 2\vert \mathcal{D}^{obs})$ | $\widehat{n_{sel}}$ | $(\hat{s}_1, \hat{s}_2\vert \widehat{n_{sel}}, \mathcal{D}^{obs})$ |
| --- | --- | --- | --- | --- | --- | --- |
| 1 | 3049–37885 (140) | 0 | 0.21 | 0.79 | 2 | (0.030, 0.023) |
| 2 | 37893–55213 (140) | 0 | 0.18 | 0.82 | 2 | (0.038, 0.027) |
| 3 | 55421–76297 (140) | 0 | 0.44 | 0.56 | 2 | (0.115, 0.014) |
| 4 | 76368–94611 (140) | 0.07 | 0.50 | 0.43 | 1 | (0.012, 0) |
| 5 | 94682–115512 (140) | 0.79 | 0.21 | 0 | 0 | (0.0, 0.0) |
| 6 | 115590–132426 (140) | 0.88 | 0.12 | 0 | 0 | (0.0, 0.0) |
| 7 | 132468–168836 (160) | 0.87 | 0.13 | 0 | 0 | (0.0, 0.0) |

**Figure 5:**
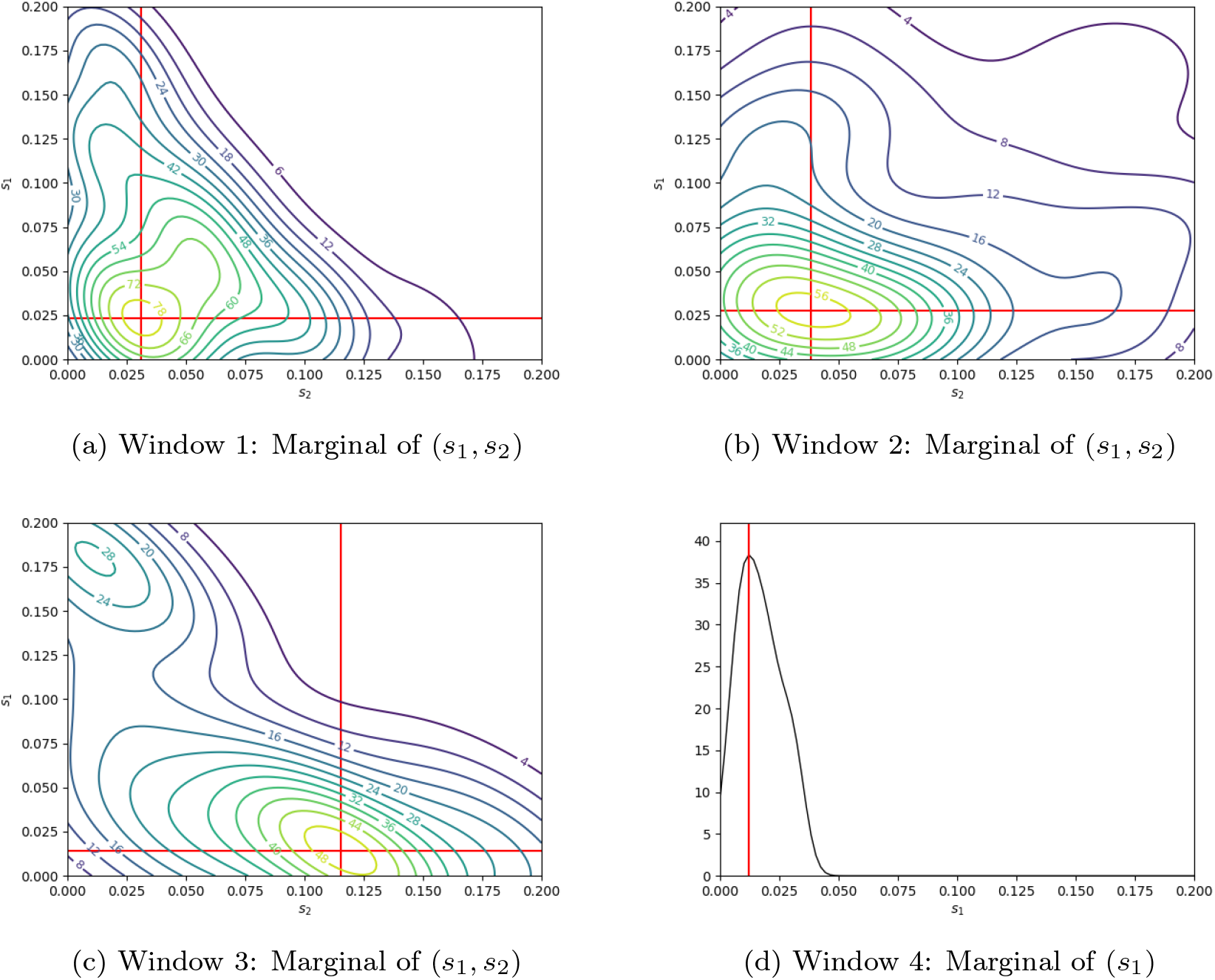
Inference on Yeast data (using 2 informative replicates). The bivariate conditional marginal posterior distributions of (*s, s*) [(a), (b), (c)] and *s*_1_ [(d)] given 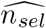 for the first 4 windows with 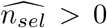 inferred using 2 replicates from the experimental data reported in Burke et al. (2014), assuming population size (*N*_*e*_ = 1000), for 7 windows on chromosome 11, each containing 140/160 SNPs.

## 6. Discussion

For experimental evolution data, methods have been proposed that test for selection. Estimates for selection coefficients *s* are also available. These methods typically operate on a SNP-by-SNP basis and do not distinguish between the selected targets and hitchhiking phenomena at neighboring positions due to linkage. Here, we propose a different window-based approach that estimates the number of selected targets and their selection coefficients. Our approach relies on high-dimensional summary statistics and a novel combination of ABC tools.

A unique feature of our method is that we also estimate the number of selected SNPs within a region of interest. As this has not been achieved by previous methods, these approaches do not provide this feature of the selected architecture. As SNP-by-SNP selection coefficient estimates are affected by linkage, they will be biased when more than one selected locus is present. It would also be unclear which of the SNP-specific estimates would carry the signal of a given selected locus with an unknown position. We believe this feature will be of interest to the field of experimental evolution, as it offers new insights into the genomic architecture of selection that were previously inaccessible.

According to our results, the estimates are fairly reliable in the haploid case if the model is correctly specified. In the diploid case, sufficiently strong selection appears to be required for reliable results.

Another advantage of our method is that it provides a posterior estimate for assessing uncertainty. Uncertainty quantification is crucial as it provides a measure of confidence in the estimated parameters.

## Acknowledgements

RD is funded by pdfRC (grant nos. EP/V025899/1, EP/T017112/1) and NERC (grant no. NE/T00973X/1). We acknowledge helpful comments by Yuexuan Wang and Marta Pelizzola.

## Supporting Information

The source code, simulated data, real data needed to reproduce the results in this article can be found at https://github.com/statrita2004/JointInference/tree/main. Additionally, the D.simulan data is available at Barghi et al. (2019)and outcrossing yeast data is available at

## Footnotes

1 𝟙(·) is used as an indicator function.

## Notes

### Competing Interest Statement

The authors have declared no competing interest.

### Summary of Updates

table 2-10 being updated and figure 5 being added to answer reviewers' concerns. Some writing were also adapted.

## References

Albert, C., Künsch, H.R., Scheidegger, A., 2015. A simulated annealing approach to approximate bayes computations. Statistics and computing 25, 1217–1232.

Barghi, N., Tobler, R., Nolte, V., Jakšić, A.M., Mallard, F., Otte, K.A., Dolezal, M., Taus, T., Kofler, R., Schlötterer, C., 2019. Genetic redundancy fuels polygenic adaptation in drosophila. PLoS biology 17, e3000128.

Beaumont, M.A., 2010a. Approximate bayesian computation in evolution and ecology. Annual review of ecology, evolution, and systematics 41, 379–406.

Beaumont, M.A., 2010b. Approximate Bayesian computation in evolution and ecology. Annual review of ecology, evolution, and systematics 41, 379–406.

Beaumont, M.A., Zhang, W., Balding, D.J., 2002. Approximate bayesian computation in population genetics. Genetics 162, 2025–2035.

Bernton, E., Jacob, P.E., Gerber, M., Robert, C.P., 2019. Approximate bayesian computation with the wasserstein distance. Journal of the Royal Statistical Society Series B: Statistical Methodology 81, 235–269.

Blum, M.G., Nunes, M.A., Prangle, D., Sisson, S.A., others, 2013. A comparative review of dimension reduction methods in approximate Bayesian computation. Statistical Science 28, 189–208.

Burke, M.K., Liti, G., Long, A.D., 2014. Standing genetic variation drives repeatable experimental evolution in outcrossing populations of saccharomyces cerevisiae. Molecular biology and evolution 31, 3228–3239.

Collin, F.d., Durif, G., Raynal, L., Lombaert, E., Gautier, M., Vitalis, R., Marin, J.M., Estoup, A., 2021. Extending approximate Bayesian computation with supervised machine learning to infer demographic history from genetic polymorphisms using DIYABC Random Forest. Molecular Ecology Resources 21, 2598–2613.

Csilléry, K., Blum, M.G., Gaggiotti, O.E., François, O., 2010. Approximate Bayesian computation (ABC) in practice. Trends in Ecology & Evolution 25, 410–418.

Dawid, A.P., 2007. The geometry of proper scoring rules. Annals of the Institute of Statistical Mathematics 59, 77–93.

DeGiorgio, M., Huber, C.D., Hubisz, M.J., Hellmann, I., Nielsen, R., 2016. Sweepfinder2: increased sensitivity, robustness and flexibility. Bioinformatics 32, 1895–1897.

Del Moral, P., Doucet, A., Jasra, A., 2012. An adaptive sequential monte carlo method for approximate bayesian computation. Statistics and computing 22, 1009–1020.

Drovandi, C.C., Pettitt, A.N., 2011. Estimation of parameters for macroparasite population evolution using approximate Bayesian computation. Biometrics 67, 225–233.

Dutta, R., Schoengens, M., Pacchiardi, L., Ummadisingu, A., Widmer, N., Künzli, P., Onnela, J.P., Mira, A., 2021. ABCpy: A high-performance computing perspective to approximate bayesian computation. Journal of Statistical Software 100, 1–38. URL: https://www.jstatsoft.org/index.php/jss/article/view/v100i07, xdoi:10.18637/jss.v100.i07.

Ewens, W.J., 2004. Mathematical population genetics: theoretical introduction. volume 1. Springer.

Fay, J.C., Wu, C.I., 2000. Hitchhiking under positive darwinian selection. Genetics 155, 1405–1413.

Fearnhead, P., Prangle, D., 2012. Constructing summary statistics for approximate Bayesian computation: semi-automatic approximate Bayesian computation [with Discussion]. Journal of the Royal Statistical Society. Series B (Statistical Methodology) 74, 419–474.

Feder, A.F., Kryazhimskiy, S., Plotkin, J.B., 2014. Identifying signatures of selection in genetic time series. Genetics 196, 509–522.

Ferrer-Admetlla, A., Liang, M., Korneliussen, T., Nielsen, R., 2014. On detecting incomplete soft or hard selective sweeps using haplotype structure. Molecular biology and evolution 31, 1275–1291.

Fisher, R.A., 1930. The genetical theory of natural selection. Oxford University Press.

Foll, M., Gaggiotti, O., 2008. A genome-scan method to identify selected loci appropriate for both dominant and codominant markers: a bayesian perspective. Genetics 180, 977–993.

Foll, M., Shim, H., Jensen, J.D., 2015. Wfabc: a w right–f isher abc-based approach for inferring effective population sizes and selection coefficients from time-sampled data. Molecular ecology resources 15, 87–98.

Fraïsse, C., Popovic, I., Mazoyer, C., Spataro, B., Delmotte, S., Romiguier, J., Loire, E., Simon, A., Galtier, N., Duret, L., et al., 2021. Dils: Demographic inferences with linked selection by using abc. Molecular Ecology Resources 21, 2629–2644.

Frazier, D.T., 2020. Robust and efficient approximate bayesian computation: A minimum distance approach. arXiv preprint 2006.14126.

Frazier, D.T., Martin, G.M., Robert, C.P., Rousseau, J., 2018. Asymptotic properties of approximate Bayesian computation. Biometrika 105, 593– 607.

Frazier, D.T., Nott, D.J., Drovandi, C., Kohn, R., 2023. Bayesian inference using synthetic likelihood: asymptotics and adjustments. Journal of the American Statistical Association 118, 2821–2832.

Fujisawa, M., Teshima, T., Sato, I., Sugiyama, M., 2021. γ-abc: Outlierrobust approximate bayesian computation based on a robust divergence estimator, in: International Conference on Artificial Intelligence and Statistics, PMLR. pp. 1783–1791.

Gneiting, T., Raftery, A.E., 2007. Strictly proper scoring rules, prediction, and estimation. Journal of the American Statistical Association 102, 359–378. doi:10.1198/016214506000001437.

Gutmann, M.U., Dutta, R., Kaski, S., Corander, J., 2018. Likelihood-free inference via classification. Statistics and Computing 28, 411–425.

Haller, B.C., Messer, P.W., 2019. SLiM 3: forward genetic simulations beyond the Wright–Fisher model. Molecular Biology and Evolution 36, 632– 637.

Hartig, F., Calabrese, J.M., Reineking, B., Wiegand, T., Huth, A., 2011. Statistical inference for stochastic simulation models–theory and application. Ecology letters 14, 816–827.

He, Z., Dai, X., Beaumont, M., Yu, F., 2020. Maximum likelihood estimation of natural selection and allele age from time series data of allele frequencies. bioRxiv, 837310.

Iranmehr, A., Akbari, A., Schlötterer, C., Bafna, V., 2017. Clear: Composition of likelihoods for evolve and resequence experiments. Genetics 206, 1011–1023.

Jiang, B., 2018. Approximate bayesian computation with kullback-leibler divergence as data discrepancy, in: International conference on artificial intelligence and statistics, PMLR. pp. 1711–1721.

Johri, P., Charlesworth, B., Jensen, J.D., 2020. Toward an evolutionarily appropriate null model: jointly inferring demography and purifying selection. Genetics 215, 173–192.

Jónás, Á., Taus, T., Kosiol, C., Schlötterer, C., Futschik, A., 2016. Estimating the effective population size from temporal allele frequency changes in experimental evolution. Genetics 204, 723–735.

Jorde, P.E., Ryman, N., 2007. Unbiased estimator for genetic drift and effective population size. Genetics 177, 927–935.

Kelly, J.K., Hughes, K.A., 2019. Pervasive linked selection and intermediatefrequency alleles are implicated in an evolve-and-resequencing experiment of drosophila simulans. Genetics 211, 943–961.

Kim, Y., Nielsen, R., 2004. Linkage disequilibrium as a signature of selective sweeps. Genetics 167, 1513–1524.

Kim, Y., Stephan, W., 2002. Detecting a local signature of genetic hitchhiking along a recombining chromosome. Genetics 160, 765–777.

Lacerda, M., Seoighe, C., 2014. Population genetics inference for longitudinally-sampled mutants under strong selection. Genetics 198, 1237–1250.

Lenormand, M., Jabot, F., Deffuant, G., 2013. Adaptive approximate bayesian computation for complex models. Computational Statistics 28, 2777–2796.

Li, W., Fearnhead, P., 2018. On the asymptotic efficiency of approximate Bayesian computation estimators. Biometrika 105, 285–299.

Lintusaari, J., Gutmann, M.U., Dutta, R., Kaski, S., Corander, J., 2017. Fundamentals and recent developments in approximate bayesian computation. Systematic biology 66, e66–e82.

Malaspinas, A.S., Malaspinas, O., Evans, S.N., Slatkin, M., 2012. Estimating allele age and selection coefficient from time-serial data. Genetics 192, 599– 607.

Nguyen, H.D., Arbel, J., Lü, H., Forbes, F., 2020. Approximate bayesian computation via the energy statistic. IEEE access 8, 131683–131698.

Nielsen, R., Williamson, S., Kim, Y., Hubisz, M.J., Clark, A.G., Bustamante, C., 2005. Genomic scans for selective sweeps using snp data. Genome research 15, 1566–1575.

Pacchiardi, L., Khoo, S., Dutta, R., 2024. Generalized bayesian likelihood-free inference. Electronic Journal of Statistics 18, 3628–3686.

Paris, C., Servin, B., Boitard, S., 2019. Inference of selection from genetic time series using various parametric approximations to the wright-fisher model. G3: Genes, Genomes, Genetics 9, 4073–4086.

Park, M., Jitkrittum, W., Sejdinovic, D., 2016. K2-abc: Approximate bayesian computation with kernel embeddings, in: Artificial intelligence and statistics, PMLR. pp. 398–407.

Pavlidis, P., Živković, D., Stamatakis, A., Alachiotis, N., 2013. Sweed: likelihood-based detection of selective sweeps in thousands of genomes. Molecular biology and evolution 30, 2224–2234.

Peng, B., Kimmel, M., 2005. simuPOP: a forward-time population genetics simulation environment. Bioinformatics 21, 3686–3687.

Pritchard, J.K., Seielstad, M.T., Perez-Lezaun, A., Feldman, M.W., 1999. Population growth of human y chromosomes: a study of y chromosome microsatellites. Molecular biology and evolution 16, 1791–1798.

Rizzo, M.L., Székely, G.J., 2016. Energy distance. wiley interdisciplinary reviews: Computational statistics 8, 27–38.

Sackman, A.M., Harris, R.B., Jensen, J.D., 2019. Inferring demography and selection in organisms characterized by skewed offspring distributions. Genetics 211, 1019–1028.

Schraiber, J.G., Evans, S.N., Slatkin, M., 2016. Bayesian inference of natural selection from allele frequency time series. Genetics 203, 493–511.

Sisson, S.A., Fan, Y., Beaumont, M., 2018. Handbook of approximate Bayesian computation. CRC Press.

Tajima, F., 1983. Evolutionary relationship of dna sequences in finite populations. Genetics 105, 437–460.

Taus, T., Futschik, A., Schlötterer, C., 2017. Quantifying selection with pool-seq time series data. Molecular biology and evolution 34, 3023–3034.

Terhorst, J., Schlötterer, C., Song, Y.S., 2015. Multi-locus analysis of genomic time series data from experimental evolution. PLoS genetics 11, e1005069.

Thornton, K.R., 2014. A C++ template library for efficient forward-time population genetic simulation of large populations. Genetics 198, 157–166.

Topa, H., Jónás, Á., Kofler, R., Kosiol, C., Honkela, A., 2015. Gaussian process test for high-throughput sequencing time series: application to experimental evolution. Bioinformatics 31, 1762–1770.

Vlachos, C., Burny, C., Pelizzola, M., Borges, R., Futschik, A., Kofler, R., Schlötterer, C., 2019. Benchmarking software tools for detecting and quantifying selection in evolve and resequencing studies. Genome biology 20, 1–11.

Vlachos, C., Kofler, R., 2018. Mimicree2: Genome-wide forward simulations of evolve and resequencing studies. PLoS computational biology 14, e1006413.

Waples, R.S., 1989. A generalized approach for estimating effective population size from temporal changes in allele frequency. Genetics 121, 379–391.

Wiberg, R.A.W., Gaggiotti, O.E., Morrissey, M.B., Ritchie, M.G., 2017. Identifying consistent allele frequency differences in studies of stratified populations. Methods in ecology and evolution 8, 1899–1909.

Wood, S.N., 2010. Statistical inference for noisy nonlinear ecological dynamic systems. Nature 466, 1102–1104.

Wright, S., 1931. Evolution in mendelian populations. Genetics 16, 97.

Yuan, X., Miller, D.J., Zhang, J., Herrington, D., Wang, Y., 2012. An overview of population genetic data simulation. Journal of Computational Biology 19, 42–54.

